# FoldToken2: Learning compact, invariant and generative protein structure language

**DOI:** 10.1101/2024.06.11.598584

**Authors:** Zhangyang Gao, Cheng Tan, Stan Z. Li

**Affiliations:** AI Lab, Research Center for Industries of the Future, Westlake University

## Abstract

The equivariant nature of 3D coordinates has posed long term challenges in protein structure representation learning, alignment, and generation. Can we create a compact and invariant language that equivalently represents protein structures? Towards this goal, we propose FoldToken2 to transfer equivariant structures into discrete tokens, while maintaining the recoverability of the original structures. From FoldToken1 to FoldToken2, we improve three key components: (1) invariant structure encoder, (2) vector-quantized compressor, and (3) equivariant structure decoder. We evaluate FoldToken2 on the protein structure reconstruction task and show that it outperforms previous FoldToken1 by 20% in TMScore and 81% in RMSD. FoldToken2 is likely the first method that works well for both single-chain and multi-chain protein structure quantization. We believe that FoldToken2 will inspire further improvement in protein structure representation, alignment, and generation tasks. Online example is available at <u>Colab</u>.

## 1 Introduction

> “SE-(3) structure should not be special and difficult. Let’s lower the barrier.”
>
> – Our Goal

Protein structure modelling plays a foundational role in computational biology and has attracted increasing attention in machine learning. Due to the SE-(3) equivariant nature, encoding [25, 8, 11, 10, 14] and generating [18, 3, 19, 22, 1] protein structure are never trivial, which requires the special design targeted to protein structures. For example, PiFold [11] proposes the invariant featurizer to encode structure patterns, and AlphaFold2 [18] design frame-based model to generate equivariant 3D coordinates. While numerious innovations have been proposed in designing the protein structure models, the structures data itself remains SE-(3) in nature. *Can we transform equivariant structures into an invariant form, and then use the existing NLP/CV models to encode and generate the structures?*

We introduce FoldToken2, a novel method to transform SE-(3) structures into invariant representations. The key insight is to create a compact invariant latent space that preserve the structure information via self-reconstruction. After pretraining, the invariant latent representation can serve as a prototype of the equivariant structures that is editable in the latent space. Akin to image and text, we also introduce a vector quantization module to discretize the latent space, creating a SE-(3) invariant language. Taking the invariant latent embedding or language as input, we can use the existing CV or NLP models [4, 7, 9, 21] to encode and generate the protein structures. FoldToken2 contains three key components: (1) invariant encoder, (2) vector quantization module, and (3) equivariant decoder.

A frame-based GNN (BlockGAT) is used for encoding equivariant structures as invariant embeddings. FoldToken1 [12, 13] represent backbone structures as bond and torsion angles, lacking the ability to capture the 3D dependencies. As a remedy, FoldToken2 represent residues as block like AlphaFold2 [18], with a efficient and powerful graph neural network BlockGAT [15]. The BlockGAT contains a simplified featurizer for capturing the informative 3D dependencies and an optimized graph network module for learning high-level representations. The sparse graph attention mechanism make it much more efficient than the SE-(3) transformers, which is important for large-scale pre-training.

An optimized vector quantization module (SoftCVQ) is used to quantize the continuous embeddings into discrete tokens, termed fold language. There has been great success in applying vector quantization to image and video modeling; however, few attempts have been made to apply it to protein structure modeling [12, 13]. To make advanced sequence models, such as BERT [6] and GPT [4], to be powerful structure learners, quantifying continuous embeddings into discrete tokens is crucial. Based on FoldToken1 [12, 13], we further improve the vector quantization module (SoftCVQ) by proposing a new teacher-guided temperature annealing strategy, leading to improved SoftCVQ. Thanks to the elaborate design, the reconstruction results consistently outperform baselines.

A conditional SE-(3) decoder is proposed for structure generation. The decoder takes discrete tokens as conditional features and generates the 3D coordinates via equivariant graph message passing. Each SE-(3) layer include a BlockGAT and a plug-and-play SE-(3) module; By stacking multiple SE-(3) BlockGATs, we iteratively refine the protein structure from Gaussian noise conditioned on discrete tokens. During reconstruction, the BlockGAT extracts pairwise residue interactions online, and SE-(3) module uses frame-level message passing to refine the local frames of residues.

We evaluate the reconstruction performance of FoldToken2 in both single-chain and multi-chain settings. In single chain reconstruction, following FoldToken1 [12, 13] which may be the first to tokenizing single-chain, FoldToken2 significantly improves the reconstruction performance on both TMScore and RMSD by 20% and 81%, respectively. In addition, FoldToken2 extends the idea to multi-chain protein structure reconstruction, and also lead to promising results. We believe that FoldToken2 will inspire further improvement in protein structure representation learning, structure alignment, and structure generation tasks.

## 2 Method

### 2.1 Overall Framework

As shown in Fig.1, the overall framework keeps the same as FoldToken1 [12, 13], including encoder, quantifier and decoder. FoldToken2 comprehensively improves each module to enhance reconstruction performance, summarized as follows::

**Figure 1:**
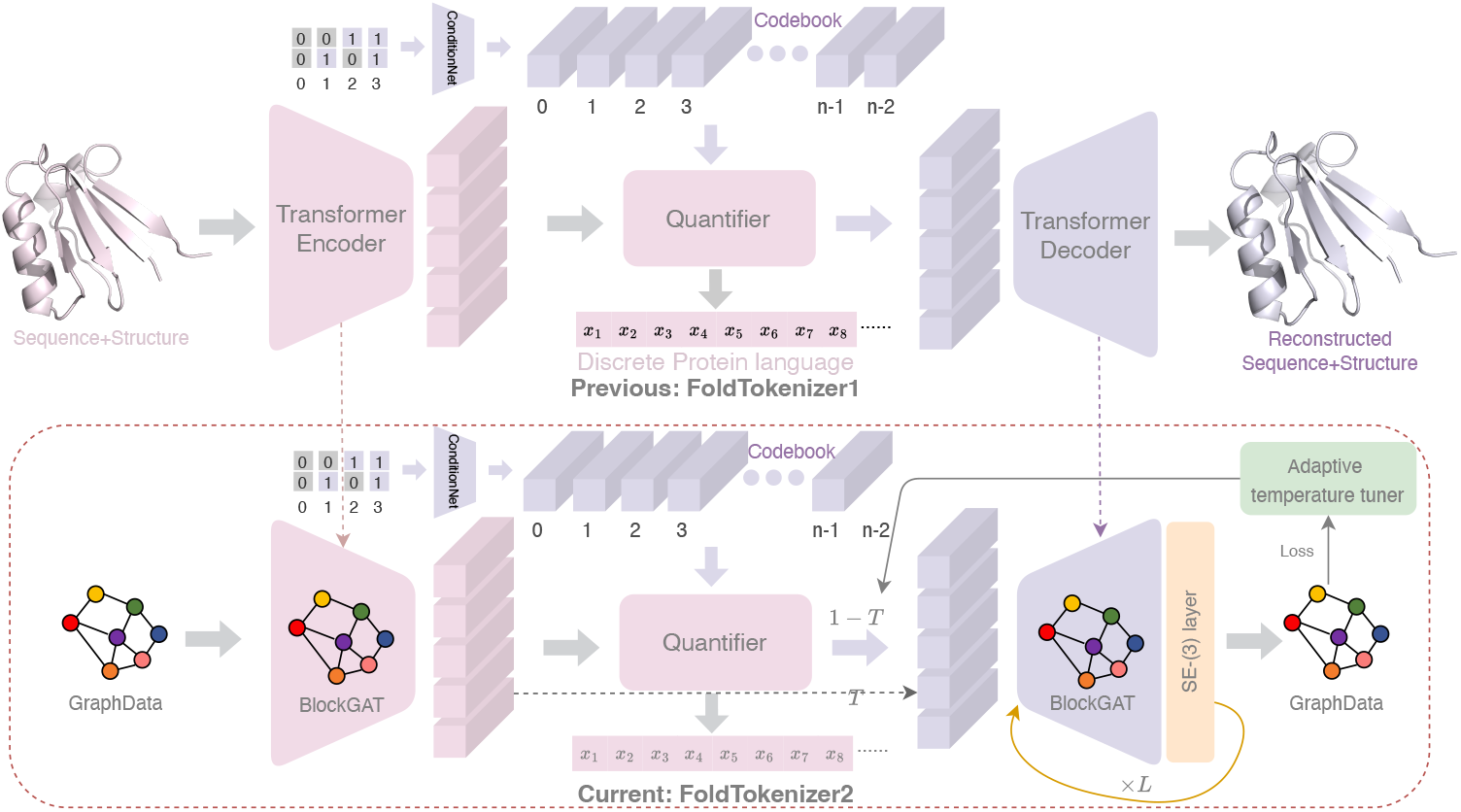
The overall framework of FoldTokenizer2, which contains contains encoder, quantifier, and decoder. In FoldToken2, we use BlockGAT to encoder protein structures as invariant embeddings, SoftCVQ to quantize the embeddings into discrete tokens, and SE-(3) layer to recover the protein structures iteratively.

**Figure 2:**
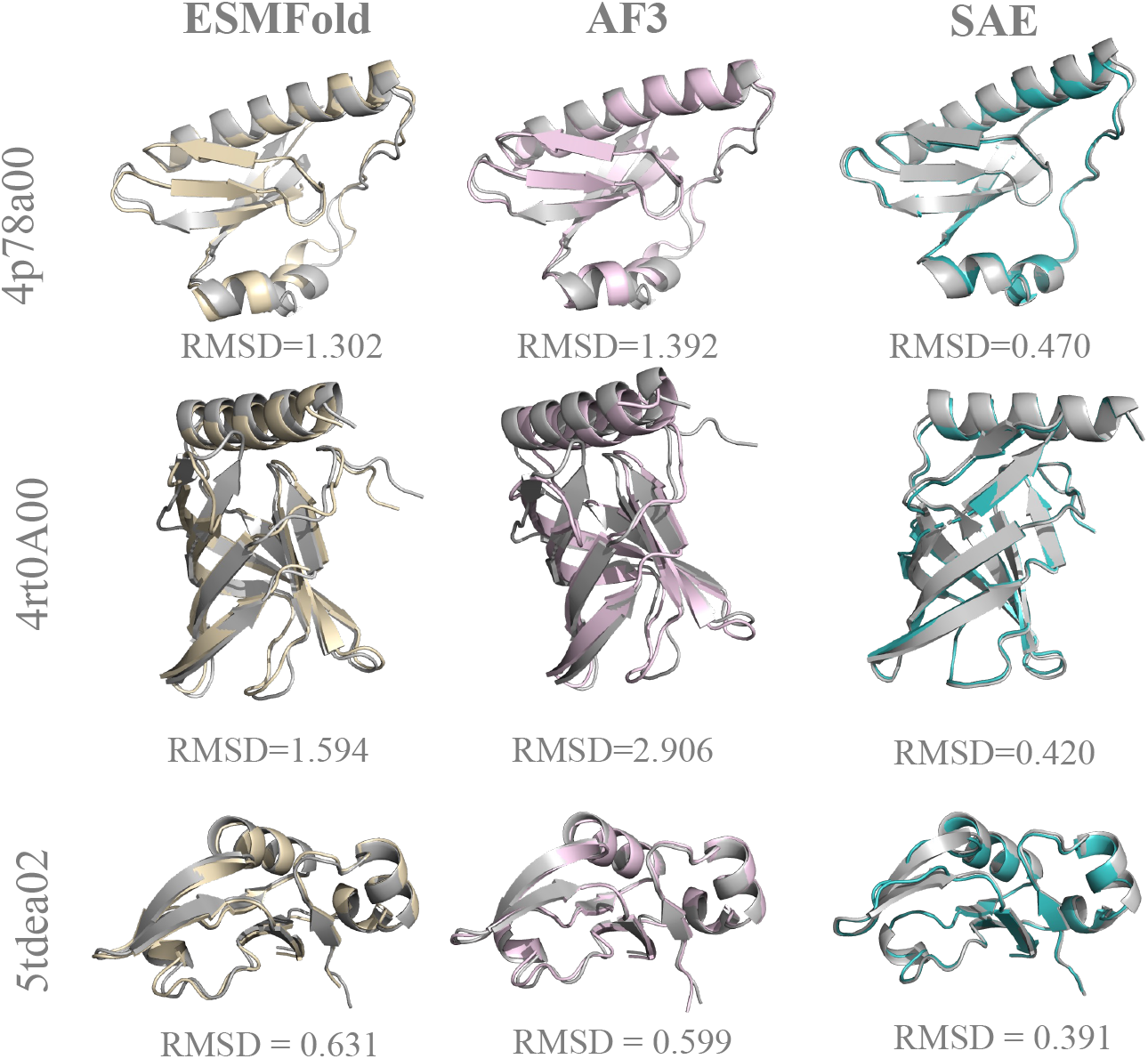
Reconstruction performance without VQ. “SAE” means structure autoencoder without VQ.

1. **Data Form**: Changing from the angle-based representation to coordinate-based one.
2. **Backbone**: We replace the transformer backbone with a new GNN, named BlockGAT.
3. **VQ**: We introduce teacher-guided temperature annealing strategy.
4. **Generator**: We propose novel SE-(3) layer to refine the protein structures iteratively.

### 2.2 Invariant Graph Encoder

Due to the rotation and translation equivariant nature, the same protein may have different coordinate records, posing a challenge in learning compact invariant representations for the same protein. Previous works [11, 17, 5, 15] have shown that the invariant featurizer can encode the invariant structure patterns, and we follow the same road: representing the protein structures as a graph consisting of invariant node and edge features. Then we use the BlockGAT [15] to learn high-level representations.

#### Frame-based Block Graph

Given a protein 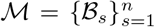 containing *n* blocks, where each block represents an amino acid, we build the block graph 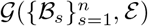 using kNN algorithm. In the block graph, the *s*-th node is represented as ℬ_*s*_ = (*T*_*s*_, ***f***_*s*_), and the edge between (*s, t*) is represented as ℬ_*st*_ = (*T*_*st*_, ***f***_*st*_). *T*_*s*_ and 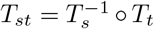 are the local frames of the *s*-th and the relative transform between the *s*-th and *t*-th blocks, respectively. ***f***_*s*_ and ***f***_*st*_ are the node and edge features.

#### BlockGAT Encoder

We use the BlockGAT [15] layer *f*_*θ*_ to learn block-level representations:

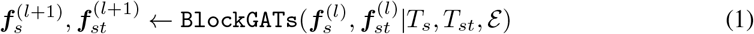

where 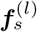 and 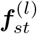 represent the input node and edge features of the *l*-th layer. *T* = (*R*, ***t***) is the local frame of the *s*-th block, and 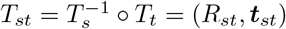 is the relative transform between the *s*-th and *t*-th blocks. 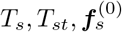 and 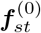 are initialized from the ground truth structures using the invariant featurizer proposed in UniIF [15].

### 2.3 Quantifier

Following FoldToken1 [12, 13], we use SoftCVQ to quantize the invariant embeddings into discrete tokens, termed fold language. Instead of projecting continuous embeddings into discrete tokens, SoftCVQ maps pre-defined binary number (***b***_*j*_) into continuous token embeddings ***v***_*j*_ and then conduct soft alignment between the token embeddings ***v***_*j*_ and latent embeddings ***h***_*s*_.

Given the decimal integer *z* and the codebook size *m*, we represent *z* as a binary vector ***b***_*j*_ with length log_2_(*m*). For example, if *m* = 4, we have ***b***_1_ = [0, 0], ***b***_2_ = [0, 1], ***b***_3_ = [1, 0], ***b***_4_ = [1, 1]. The quantization operation transforms continuous embeddings ***h***_*s*_ into discrete tokens *z*:

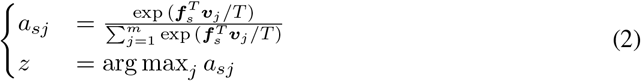

where ***v***_*j*_ is the *j*-th token embedding, generated by a conditional network, which also server as the de-quantization operation:

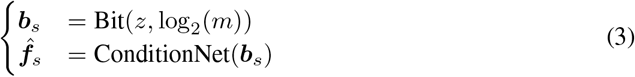

In Eq. 2, the “argmax” is non-differentiable, and we use a soft approximation during training:

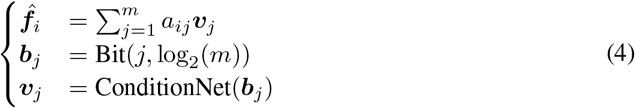

The temperature parameter *T* controls the softness of the attention query operation. When *T* is large, the attention weights will be close to uniform; otherwise, the attention weights will be close to one-hot.

During training, the temperature is reduced from 1.0 to 1e-8. The ConditionNet : 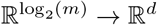 is a MLP. If the codebook size is 2^16^, the MLP projects 65536 16-dimension boolvectors into 65536 *d*-dimension vectors. Considering the gradient of the softmax operation:

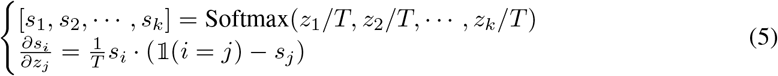

when *T* tend to zero, the gradient explodes. To avoid this, we frozen the encoder and VQ module when the temperature is lower than 1e-6. The temperature scheduler is:

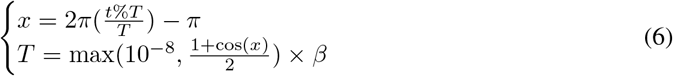

To accelerate model convergence, we introduce:

- **Teacher-Guided VQ**: We randomly copy the encoder embedding ***f***_*s*_ as the quantized feature 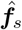 in a probability of *T*, termed teacher-guided vector quantization.
- **Adaptive tuner**: We adaptively adjust the temperature scale *β* based on training loss. The adaptive temperature tuner is defined as:

The adaptive temperature tuner is defined as:

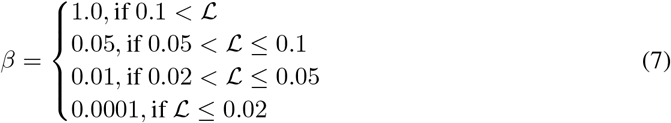

### 2.4 Equivariant Graph Decoder

Generating the protein structures conditioned on invariant representations poses significant challenges in computing efficiency. For example, training well-known AlphaFold2 from scratch takes 128 TPUv3 cores for 11 days [24]; OpenFold takes 50000 GPU hours for training [2]. In this work, we propose an efficiency plug-and-play SE(3)-layer that could be added to any GNN layer for structure prediction. Thanks to the simplified module of the SE(3)-layerand BlockGAT with sparse graph attention, we can train the model on the entire PDB dataset in 1 day using 8 NVIDIA-A100s.

#### SE-(3) Frame Passing Layer

We introduce frame-level message passing, which updates the local frame of the *s*-th block by aggregating the relative rotation *R*_*s*_ and translation ***t***_*s*_ from its neighbors:

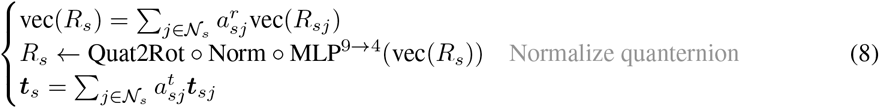

where 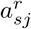 and 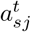 are the rotation and translation weights, and 𝒩_*s*_ is the neighbors of the *s*-th block. vec(*·*) flattens 3 *×* 3 matrix to 9-dimensional vector. MLP^9→4^(*·*) maps the 9-dim rotation matrix to 4-dim quaternion, and Norm(*·*) normalize the quaternion to ensure it represents a valid rotation. Quat2Rot(*·*) is the quaternion to rotation function. We further introduce the details as follows:

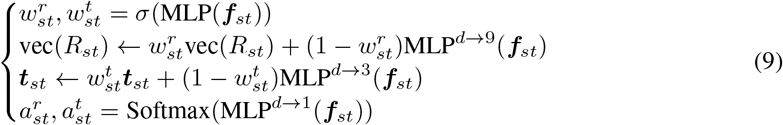

where 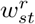 and 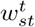 are the updating weights for rotation and translation, 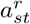 and 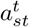 are the attention weights. The propose SE-(3) layer could be add to any graph neural network for local frame updating.

#### Iterative Refinement

We propose a new module named SE-(3) BlockGAT by adding a SE-(3) layer to BlockGAT. We stack multi-layer SE-(3) BlockGAT to iteratively refine the structures:

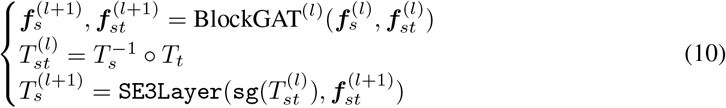

where `sg`(*·*) is the stop-gradient operation, and `SE3Layer`(*·*) is the SE-(3) layer described in Eq.9. Given the predicted local frame 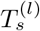, we can obtain the 3D coordinates by:

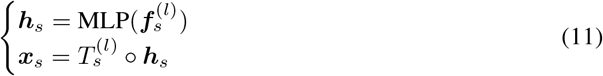

### 2.5 Reconstruction Loss

Inspired by Chroma [16], we use multiple loss functions to train the model. The overall loss is:

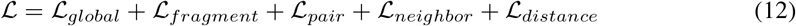

To illustrate the loss terms, we define the aligned RMSD loss as 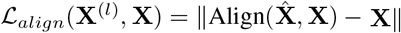, given the the ground truth 3D coordinates **X?**∈ ℝ ℝ ^*n*,3^ and the predicted 3D coordinates 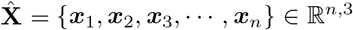. The global, fragment and pair loss are defined by the aligned MSE loss, but with different input data shape:

- **Global Loss**: **X** with shape [*n*, 4, 3]. RMSD of the global structure.
- **Fragment Loss**: **X** with shape [*n, c*, 4, 3]. RMSD of *c* neighbors for each residue.
- **Pair Loss**: **X** with shape [*n, K, c ·* 2, 4, 3]. RMSD of *c* neighbors for each kNN pair.
- **Neighbor Loss**: **X** with shape [*n, K*, 4, 3]. RMSD of *K* neighbors for each residue.

where *n* is the number of residues, *c* = 7 is the number of fragments, *K* = 30 is the number of kNN, 4 means we consider four backbone atoms *{N, CA, C, O}*, and 3 means the 3D coordinates. The distance loss is defined as the MSE loss between the predicted and ground truth pairwise distances:

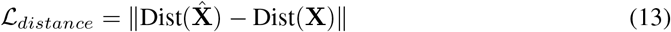

where Dist(**X**) ∈ ℝ ^*n*,*n*^ is the pairwise distance matrix of the 3D coordinates **X**. We apply the loss on each decoder layer, and the final loss is the average, whcih is crucial for good performance.

## 3 Experiments

We conduct systematic experiments to inspire further improvement in FoldToken2.

- **AutoEncoder Training (Q1):** How well the compact and invariant latent space learned by FoldToken2 can be used to reconstruct protein structures?
- **Single-Chain Reconstruction (Q2):** Can FoldToken2 outperform FoldToken1 in single-chain protein reconstruction?
- **Multi-Chain Reconstruction (Q3):** Can FoldToken2 perform well on multi-chain protein reconstruction?

### Metrics

We evaluate FoldToken2 on the protein structure reconstruction task, where the TMscore and aligned RMSD are reported.

### Single-chain Data

We use the CATH4.3 dataset to train the single-chain model. The same as protein inverse folding models, CATH4.3 is split into training (16631), validation (1516), and testing (1864) sets according to the CAT code. During evaluation, we remove proteins with NaN coordinates, resulting in 493 core samples in the final testing set (T493). Due to the different data representations, backbone architecture and iterative refinement strategies, FoldToken2 trains much slower than FoldToken1. Therefore, we do not train FoldToken2 in the AF2DB dataset like FoldToken1, which can be done in the future if the computing resource is enough. In addition, we also report results over the sub-testing set (T116) of FoldToken1 for head-to-head comparision, which contains 116 proteins.

### Multi-chain Data

We train the model using all proteins collected from the PDB dataset as of 1 March 2024. After filtering residues with missing coordinates and proteins less than 30 residues, we obtain 162K proteins for training. We random crop long proteins to ensure that the maximum length is 500. Protein complexes are supported by adding chain encoding features *c*_*ij*_ to the edge *e*_*ij*_: *c*_*ij*_ = 0, if *i* and *j* are in different chains; else *c*_*ij*_ = 1.

### 3.1 AutoEncoder Training (Q1)

#### Setting

We train FoldToken2 without vector quantization to show how well the encoder-decoder can learn and generate protein structure patterns. The results could be used as the ceiling performance of FoldToken series models. We train the model up to 25 epochs with a batch size of 8 and a learning rate of 0.001. The overall model contains 8 encoding layers, 8 decoding layers, and a 128 hidden dimension, total 9.2M parameters. The training process could be done in 1 days using 1 A100 GPU. We also report the structure prediction results of ESMFold [19], AlphaFold2 [18] and AlphaFold3 [1] for evaluating the structure generation capability of FoldToken2.

#### Results

In Table. 1, we show the reconstruction results of FoldToken2. In both T493 and T116, FoldToken2 achieves good performance in TMScore and RMSD. When compared to ESMFold, AlphaFold2, and AlphaFold3, FoldToken2 can generate more accurate protein structures from invariant latent embeddings, showing the capability of the equivariant decoder. Notabely, the protein folding decoder only contains 4.9M parameters, which is much smaller than the 688.55M parameters of ESMFold’s Folding Trunk. FoldToken2 may provide new insights for the protein folding problem.

**Table 1:** AutoEncoder reconstruction results of FoldToken2.

|  | Data | TMScore |  |  | RMSD |  |  |
| --- | --- | --- | --- | --- | --- | --- | --- |
|  |  | Avg | Max | Min | Avg | Max | Min |
| FoldToken2 | T493 | 0.98 | 0.99 | 0.93 | 0.46 | 2.12 | 0.26 |
| ESMFold | T116 | 0.94 | 0.99 | 0.88 | 1.21 | 5.33 | 0.33 |
| AlphaFold2 (Colab) | T116 | 0.92 | 0.99 | 0.35 | 1.44 | 11.00 | 0.38 |
| AlphaFold3 | T116 | 0.94 | 0.99 | 0.74 | 1.24 | 4.02 | 0.33 |
| FoldToken2 | T116 | **0.98** | **0.99** | **0.87** | **0.41** | **0.87** | **0.25** |

### 3.2 Single-Chain Reconstruction (Q2)

#### Setting

We compare FoldToken2 with FoldToken1 on both T116 and T493 datasets. The model is trained up to 25 epochs with a batch size of 8 and leading rate of 0.001. One experiment could be done about 1 day using 1 80G A100 GPU.

In Table. 2 and Fig. 3, we show the reconstruction results of FoldToken2 and conclude that:

**Table 2:**
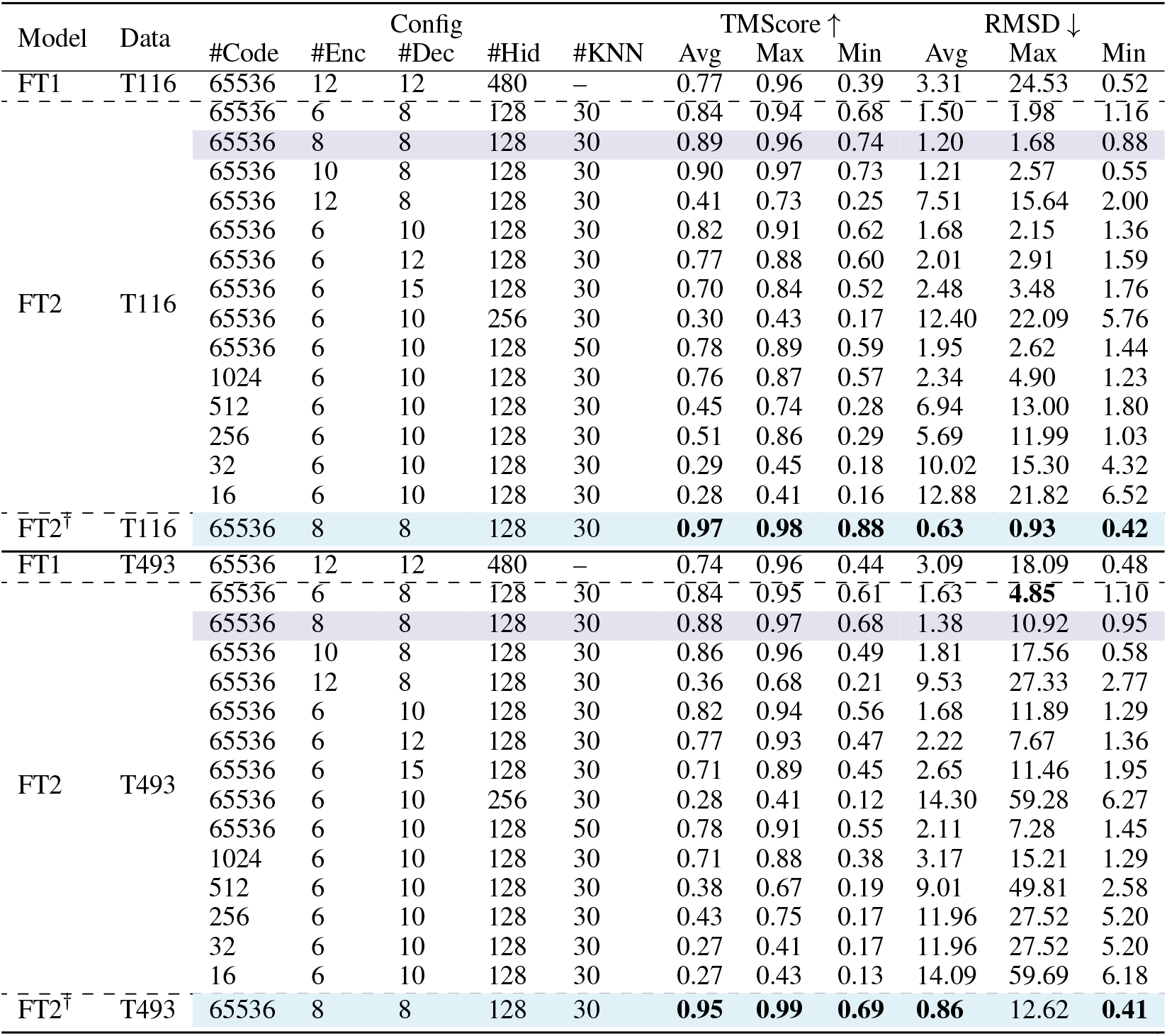
Reconstruction performance. We highlight the promising models in color. FT1, FT2, and FT2^*†*^ indicates FoldToken1 [12, 13], FoldToken2, and FoldToken2 with the adaptive temperature annealing strategy, respectively. FT2 use the linear temperature decay scheduler, reducing the temperature from 1.0 to 1*e*^−5^ in 40k steps. FT2^*†*^ use the adaptive temperature tuner, reducing the temperature from 1.0 to 1*e*^−5^ in 5k steps.

**Figure 3:**
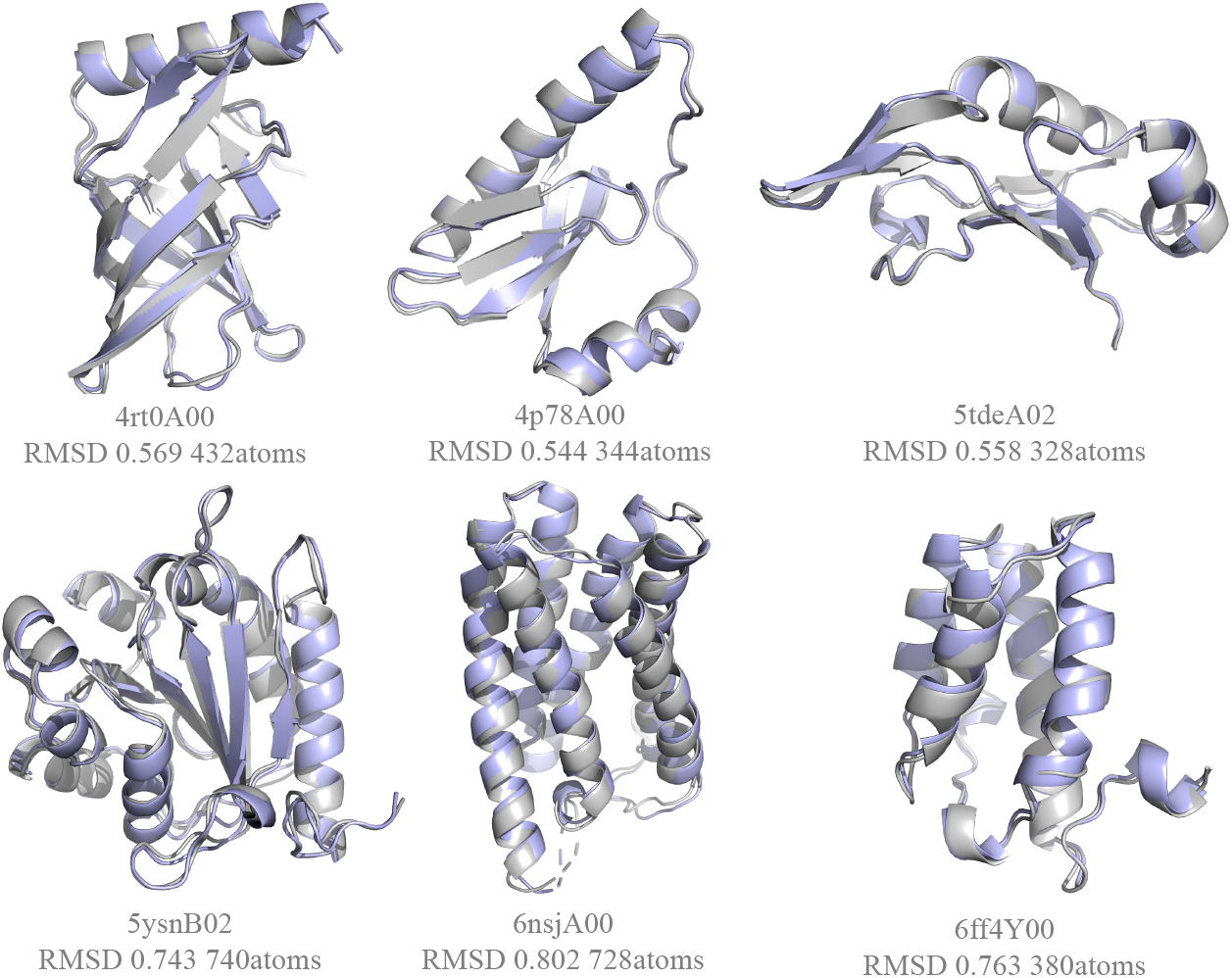
Single-chain reconstruction. Grey and colored residues represent the ground truth and predicted ones.

#### FoldToken2 outperforms FoldToken1 by a large margin in structure reconstruction

In average TMScore, the FT2^*†*^ model achieves 0.97 on T116, outperforming FT1 by 20%. In RMSD, FT2^*†*^ achieves 0.63 on T116, outperforming FoldToken1 by 81%. Similar results are observed on T493. Moreover, FT2^*†*^ is more reliable than others, with worst-case TMScores and RMSD of 0.88 and 0.42, respectively, compared to 0.39 and 24.53 for FoldToken1. These results suggest that the new encoder, quantifier, and decoder in FoldToken2 significantly enhance protein structure reconstruction.

#### Scalling up model does not perform better on limited data

We find that the 8-layer encoder and decoder with hidden size 128 achieve the best performance. Futher increasing the number of layers or hidden size does not lead to improvement, possiblly limited to the small training data volumn. Also, we should point out that we do not pay much attention to stablize the training of deepper and wider models, and results may improve with careful adjustments.

#### Enlarging Decoing KNN do not improve performance

Increasing the number of KNN neighbors in the decoder from 30 to 50 does not improve the performance. This result suggests that the 30 kNN neighbors are sufficient to capture the structure interactions, when using BlockGAT as backbone. The reason may be that the BlockGAT introduces virtual frames for considering long-term dependencies.

#### Larger codebook size leads to better reconstruction performance

Increasing the codebook size from 16 to 65,536 consistently improves reconstruction performance. This suggests that the 20-codebook size in FoldSeek is likely insufficient for accurately describing protein structures from a reconstruction perspective. However, since FoldSeek [23] focuses on structure alignment, this loss of information does not significantly affect its widespread usage.

## 4 Multi-Chain Reconstruction (Q3)

### Setting

We train FoldToken2 on the multi-chain protein reconstruction task using the PDB dataset. The model is trained for up to 25 epochs with a batch size of 8, a learning rate of 0.001, and a padding length of 500. There is no available benchmark for evaluating multi-chain protein reconstruction, and we provide some visual examples in Fig. 4. The comprehensive benchmark is coming soon.

**Figure 4:**
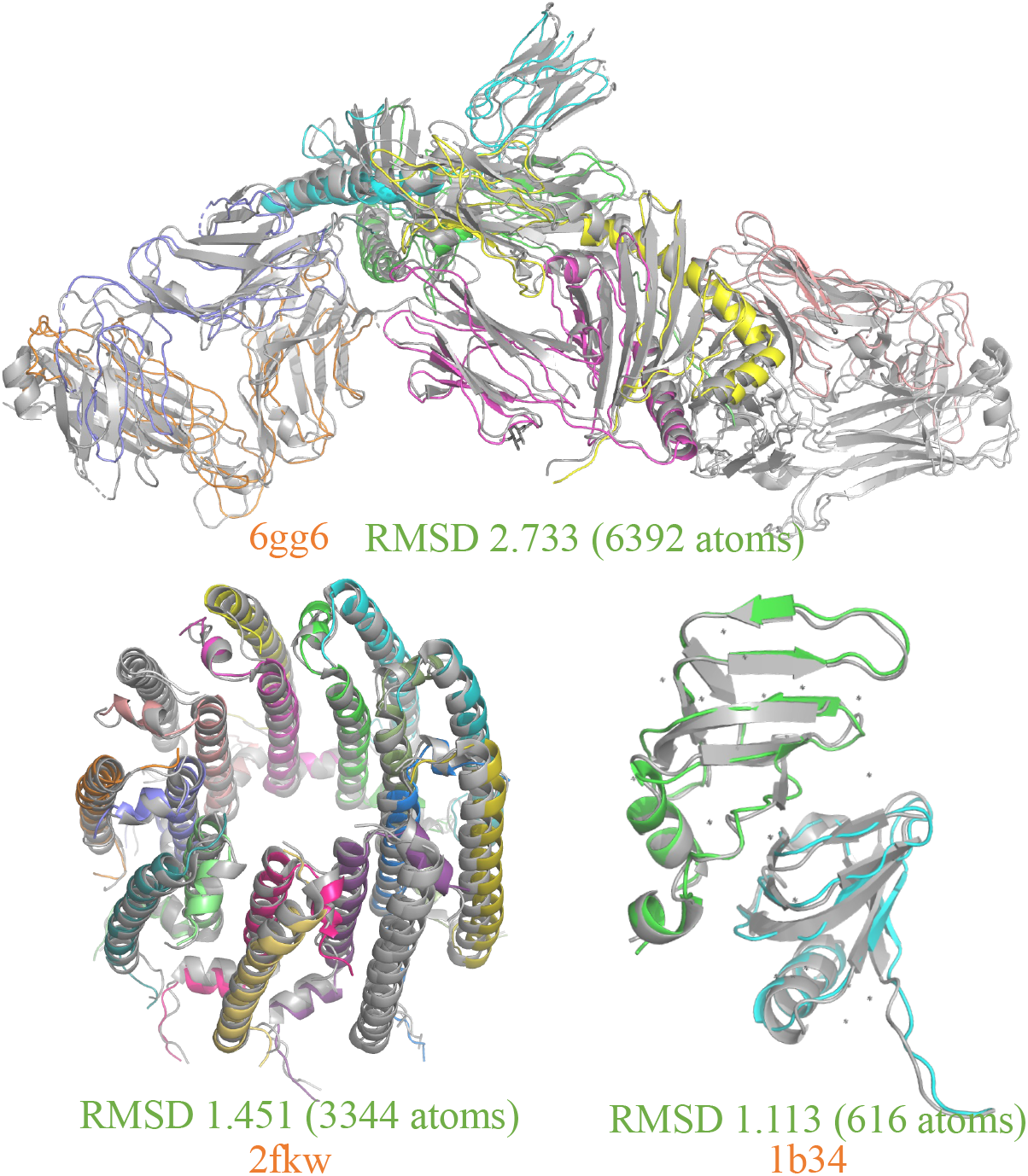
Multi-chain reconstruction. Grey and colored residues represent the ground truth and predicted ones.

**Figure 5:**
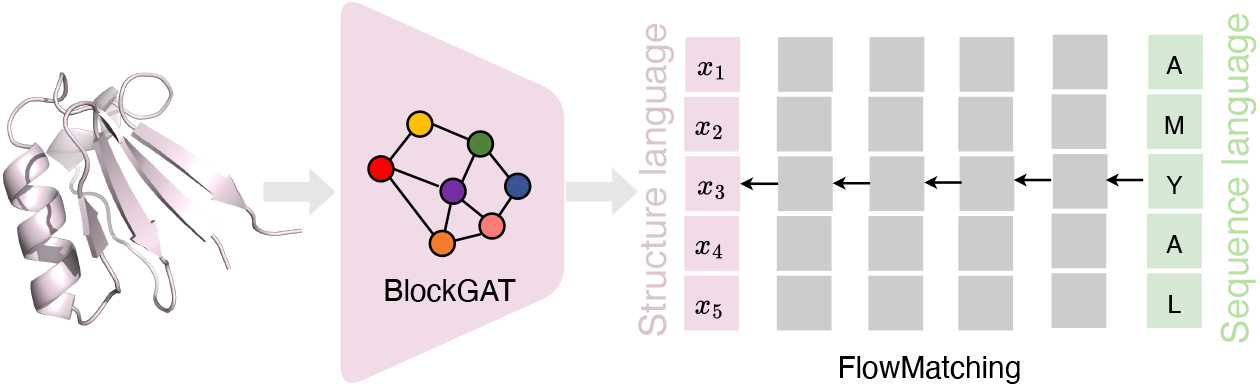
Flow matching for sequence-srtructure translation.

**Figure 6:**
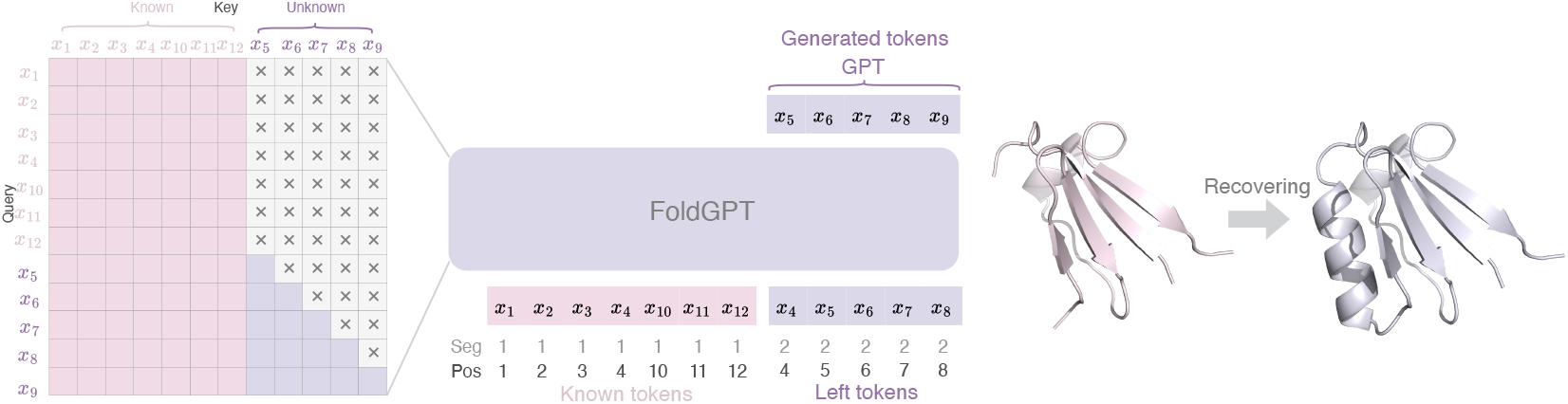
FoldGPT for srtructure generation.

### FoldToken2 generalize well to large-scale protein systems

While we crop long proteins to ensure that the maximum length is 500, FoldToken2 can generate protein complexes with low RMSD errors, as shown in Fig. 4. The results suggest that FoldToken2 can be used for multiple real-world applications beyond protein monomers.

### We are surprised by the small model size and good training efficiency

Unlike other models that use AF2 for structure generation, FoldToken2 does not copy AF2. Instead, we use a light-weight model, comprising a 4.31M encoder, 4.27M quantifier, and 4.92M decoder, achieving state-of-the-art results. When training a protein structure generation model, OpenFold requires 5000 GPU hours; in contrast, FoldToken2 only requires 40GPU hours to train on the entire PDB dataset.

## 5 Extension (Future Work)

### TokenFlow

Given a discrete protein language and protein sequence. we train a flow-matching model using rectified flow [20] on the CATH4.3 training set. The flow-matching model is designed to predict protein structures from single protein sequences without using MSA information. The model is trained for up to 1000 epochs with a batch size of 128, a learning rate of 0.0005, and a padding length of 512. The overall model architecture consists of a 12-layer transformer, similar to ESM-35M. The training process can be completed in about one day using a single A100 GPU. When training over 16K data of CATH4.3, the model seems to be not work well as expected. Limited by energy and computing resources, we plan to scale the model in the future.

### FoldGPT

Similar to FoldToken1, we train a model for generating masked structure regions based on unmasked ones. However, the model seems to be overfitting on the training data, and we need to scale up the model in the future.

## 6 Conclusion

This paper proposes FoldToken2, probably the first multi-chain protein structure tokenization approach. With great efforts in improving the encoder, quantifier, and decoder, FoldToken2 significantly outperforms FoldToken1 by 20% in TMScore and 81% in RMSD. Additionally, the training efficiency is remarkable, requiring only 40 GPU hours to train on the entire PDB dataset. The propose methods may inspire further improvement in protein structure representation learning, structure alignment, and structure generation tasks.

## References

[1] Josh Abramson, Jonas Adler, Jack Dunger, Richard Evans, Tim Green, Alexander Pritzel, Olaf Ronneberger, Lindsay Willmore, Andrew J Ballard, Joshua Bambrick, et al. Accurate structure prediction of biomolecular interactions with alphafold 3. Nature, pages 1–3, 2024.

[2] Gustaf Ahdritz, Nazim Bouatta, Christina Floristean, Sachin Kadyan, Qinghui Xia, William Gerecke, Timothy J O’Donnell, Daniel Berenberg, Ian Fisk, Niccolò Zanichelli et al. Open-fold: Retraining alphafold2 yields new insights into its learning mechanisms and capacity for generalization. Nature Methods, pages 1–11, 2024.

[3] Minkyung Baek, Frank DiMaio, Ivan Anishchenko, Justas Dauparas, Sergey Ovchinnikov, Gyu Rie Lee, Jue Wang, Qian Cong, Lisa N Kinch, R Dustin Schaeffer, et al. Accurate prediction of protein structures and interactions using a three-track neural network. Science, 373(6557):871–876, 2021.

[4] Tom Brown, Benjamin Mann, Nick Ryder, Melanie Subbiah, Jared D Kaplan, Prafulla Dhariwal, Arvind Neelakantan, Pranav Shyam, Girish Sastry, Amanda Askell, et al. Language models are few-shot learners. Advances in neural information processing systems, 33:1877–1901, 2020.

[5] Justas Dauparas, Ivan Anishchenko, Nathaniel Bennett, Hua Bai, Robert J Ragotte, Lukas F Milles, Basile IM Wicky, Alexis Courbet, Rob J de Haas, Neville Bethel, et al. Robust deep learning–based protein sequence design using proteinmpnn. Science, 378(6615):49–56, 2022.

[6] Jacob Devlin, Ming-Wei Chang, Kenton Lee, and Kristina Toutanova. Bert: Pre-training of deep bidirectional transformers for language understanding. arXiv preprint arXiv:1810.04805, 2018.

[7] Alexey Dosovitskiy, Lucas Beyer, Alexander Kolesnikov, Dirk Weissenborn, Xiaohua Zhai, Thomas Unterthiner, Mostafa Dehghani, Matthias Minderer, Georg Heigold, Sylvain Gelly, et al. An image is worth 16x16 words: Transformers for image recognition at scale. arXiv preprint arXiv:2010.11929, 2020.

[8] Hehe Fan, Zhangyang Wang, Yi Yang, and Mohan Kankanhalli. Continuous-discrete convolution for geometry-sequence modeling in proteins. In The Eleventh International Conference on Learning Representations, 2022.

[9] Zhangyang Gao, Daize Dong, Cheng Tan, Jun Xia, Bozhen Hu, and Stan Z Li. A graph is worth k words: Euclideanizing graph using pure transformer. arXiv preprint arXiv:2402.02464, 2024.

[10] Zhangyang Gao, Cheng Tan, Xingran Chen, Yijie Zhang, Jun Xia, Siyuan Li, and Stan Z Li. Kw-design: Pushing the limit of protein deign via knowledge refinement. In The Twelfth International Conference on Learning Representations, 2023.

[11] Zhangyang Gao, Cheng Tan, and Stan Z Li. Pifold: Toward effective and efficient protein inverse folding. In The Eleventh International Conference on Learning Representations, 2022.

[12] Zhangyang Gao, Cheng Tan, and Stan Z Li. Vqpl: Vector quantized protein language. arXiv preprint arXiv:2310.04985, 2023.

[13] Zhangyang Gao, Cheng Tan, Jue Wang, Yufei Huang, Lirong Wu, and Stan Z Li. Foldtoken: Learning protein language via vector quantization and beyond. arXiv preprint arXiv:2403.09673, 2024.

[14] Zhangyang Gao, Cheng Tan, Yijie Zhang, Xingran Chen, Lirong Wu, and Stan Z Li. Proteininvbench: Benchmarking protein inverse folding on diverse tasks, models, and metrics. Advances in Neural Information Processing Systems, 36, 2024.

[15] Zhangyang Gao, Jue Wang, Cheng Tan, Lirong Wu, Yufei Huang, Siyuan Li, Zhirui Ye, and Stan Z Li. Uniif: Unified molecule inverse folding. arXiv preprint arXiv:2405.18968, 2024.

[16] John B Ingraham, Max Baranov, Zak Costello, Karl W Barber, Wujie Wang, Ahmed Ismail, Vincent Frappier, Dana M Lord, Christopher Ng-Thow-Hing, Erik R Van Vlack, et al. Illuminating protein space with a programmable generative model. Nature, 623(7989):1070–1078, 2023.

[17] Bowen Jing, Stephan Eismann, Patricia Suriana, Raphael John Lamarre Townshend, and Ron Dror. Learning from protein structure with geometric vector perceptrons. In International Conference on Learning Representations, 2020.

[18] John Jumper, Richard Evans, Alexander Pritzel, Tim Green, Michael Figurnov, Olaf Ronneberger, Kathryn Tunyasuvunakool, Russ Bates, Augustin Žídek, Anna Potapenko, et al. Highly accurate protein structure prediction with alphafold. Nature, 596(7873):583–589, 2021.

[19] Zeming Lin, Halil Akin, Roshan Rao, Brian Hie, Zhongkai Zhu, Wenting Lu, Nikita Smetanin, Robert Verkuil, Ori Kabeli, Yaniv Shmueli, et al. Evolutionary-scale prediction of atomic-level protein structure with a language model. Science, 379(6637):1123–1130, 2023.

[20] Xingchao Liu, Chengyue Gong, and Qiang Liu. Flow straight and fast: Learning to generate and transfer data with rectified flow. arXiv preprint arXiv:2209.03003, 2022.

[21] Robin Rombach, Andreas Blattmann, Dominik Lorenz, Patrick Esser, and Björn Ommer. High-resolution image synthesis with latent diffusion models. In Proceedings of the IEEE/CVF conference on computer vision and pattern recognition, pages 10684–10695, 2022.

[22] Cheng Tan, Zhangyang Gao, Lirong Wu, Jun Xia, Jiangbin Zheng, Xihong Yang, Yue Liu, Bozhen Hu, and Stan Z Li. Cross-gate mlp with protein complex invariant embedding is a one-shot antibody designer. In Proceedings of the AAAI Conference on Artificial Intelligence, volume 38, pages 15222–15230, 2024.

[23] Michel Van Kempen, Stephanie S Kim, Charlotte Tumescheit, Milot Mirdita, Jeongjae Lee, Cameron LM Gilchrist, Johannes Söding, and Martin Steinegger. Fast and accurate protein structure search with foldseek. Nature biotechnology, 42(2):243–246, 2024.

[24] Guoxia Wang, Xiaomin Fang, Zhihua Wu, Yiqun Liu, Yang Xue, Yingfei Xiang, Dianhai Yu, Fan Wang, and Yanjun Ma. Helixfold: An efficient implementation of alphafold2 using paddlepaddle. arXiv preprint arXiv:2207.05477, 2022.

[25] Zuobai Zhang, Minghao Xu, Arian Jamasb, Vijil Chenthamarakshan, Aurelie Lozano, Payel Das, and Jian Tang. Protein representation learning by geometric structure pretraining. arXiv preprint arXiv:2203.06125, 2022.

